# Concentration and quantification of *Tilapia tilapinevirus* from water using a simple iron flocculation coupled with probe-based RT-qPCR

**DOI:** 10.1101/2021.08.10.455809

**Authors:** Suwimon Taengphu, Pattanapon Kayansamruaj, Yasuhiko Kawato, Jerome Delamare-Deboutteville, Chadag Vishnumurthy Mohan, Ha Thanh Dong, Saengchan Senapin

## Abstract

*Tilapia tilapinevirus* (also known as tilapia lake virus, TiLV) is an important virus responsible for die-off of farmed tilapia globally. Detection and quantification of the virus from environmental DNA/RNA (eDNA/eRNA) using pond water represents a potential, noninvasive routine approach for pathogen monitoring and early disease forecasting in aquaculture systems. Here, we report a simple iron flocculation method for viral concentration from water combined with a newly developed hydrolysis probe quantitative RT-qPCR method for detection and quantification of TiLV. The RT-qPCR method targeting a conserved region of TiLV genome segment 9 has a detection limit of 10 viral copies per µL of template. The method had a 100% analytical specificity and sensitivity for TiLV. The optimized iron flocculation method was able to recover 16.11 ± 3.3% of virus from water samples spiked with viral cultures. During disease outbreak cases from an open-caged system and a closed hatchery system, both tilapia and water samples were collected for detection and quantification of TiLV. The results revealed that TiLV was detected from both clinically sick fish and asymptomatic fish. Most importantly, the virus was successfully detected from water samples collected from different locations in the affected farms e.g. river water samples from affected cages (8.50 × 10^2^ to 2.79 × 10^4^ copies/L) and fish-rearing water samples, sewage, and reservoir (4.29 × 10^2^ to 3.53 × 10^3^ copies/L) from affected and unaffected ponds of the hatchery. In summary, this study suggests that the eRNA detection system using iron flocculation coupled with probe based-RT-qPCR is feasible for concentration and quantification of TiLV from water. This approach might be useful for noninvasive monitoring of TiLV in tilapia aquaculture systems and facilitating appropriate decisions on biosecurity interventions needed.

## Introduction

*Tilapia tilapinevirus* (commonly called tilapia lake virus, TiLV) is a novel and only virus in a new genus *Tilapinevirus* under the family *Amnoonviridae* (International Committee on Taxonomy of Viruses. 2019). Since its first discovery in 2014, the virus had significant impacts on tilapia aquaculture worldwide (Eyngor et al. 2014; Ferguson et al. 2014; Jansen et al. 2019). TiLV is an RNA virus with a 10 segmented negative sense single stranded genome of approximately 10.323 kb in size (Bacharach et al. 2016). Disease caused by TiLV usually results in cumulative mortality from 20 to 90% (Behera et al. 2018; Dong et al. 2017a; Eyngor et al. 2014; Ferguson et al. 2014; Surachetpong et al. 2017). So far, there are 16 countries that reportedly confirmed detection of TiLV (Jansen et al. 2019; Surachetpong et al. 2020), but a wider geographical spread has been hypothesized due to active movements of live tilapia with other countries (Dong et al. 2017b). Waterborne spread of TiLV might also contribute to pathogen dissemination to new areas as well as transmission to other fish species (Chiamkunakorn et al. 2019; Eyngor et al. 2014; Jaemwimol et al. 2018; Piamsomboon & Wongtavatchai 2021). Experimental evidences have already demonstrated that TiLV is both horizontally and vertically transmitted (Dong et al. 2020; Eyngor et al. 2014; Jaemwimol et al. 2018; Yamkasem et al. 2019).

With respect to waterborne transmission of fish pathogens, several studies employed various viral concentration methods from water for pathogen detection (For example, Haramoto et al. (2007); Kawato et al. (2016); Minamoto et al. (2009); Nishi et al. (2016)). The concept is one of the applications of environmental DNA (eDNA) which is nucleic acids extracted from environmental samples such as water, soil, and feces (Bass et al. 2015; Gomes et al. 2017). The eDNA gives advantages in disease monitoring, control measure design, risk factor analysis and studies of viral survival nature (example review in Oidtmann et al. (2018)). The work described by Kawato et al. (2016) used an iron flocculation method to concentrate red sea bream iridovirus (RSIV) in a challenge model with Japanese amberjack (*Seriola quinqueradiata*). Results from that study showed that detection by qPCR of RSIV from fish-rearing water samples peaked more than five days before fish mortality occurred, suggesting potential benefit of using iron flocculation method for disease forecast. Others studies used a cation□coated filter method to detect DNAs of cyprinid herpesvirus 3 (CyHV-3) (also known as koi herpesvirus, KHV) from concentrated river water samples three to four months before mass mortalities events occurred in wild carp in Japan (Haramoto et al. 2007; Minamoto et al. 2009). Additionally, the virus was still detectable in river water for at least three months after the outbreaks (Minamoto et al. 2009). These findings helped local authorities and farmers to make rapid decisions for emergency harvest, biosecurity implementation, follow appropriate disinfection procedures and fallowing periods.

Several molecular methods have been developed for detection of TiLV including RT-PCR (Eyngor et al. 2014), nested and semi-nested PCR (Dong et al. 2017a; Kembou Tsofack et al. 2017; Taengphu et al. 2020), RT-qPCR (Tattiyapong et al. 2018; Waiyamitra et al. 2018), loop-mediated isothermal amplification (LAMP) (Kampeera et al. 2021; Phusantisampan et al. 2019; Yin et al. 2019) and Nanopore-based PCR amplicon approach (Delamare-Deboutteville et al. 2021). However, all of these methods target fish tissue specimens for diagnosis, none of which reported any application for TiLV detection from environmental water samples. Previous probe-based RT-qPCR methods developed to detect TiLV from tilapia clinical samples with detection limits of 2.7×10^4^ or ∼70,000 copies (Kembou Tsofack et al. 2017; Waiyamitra et al. 2018) might not be sensitive enough to detect low viral loads of TiLV in environmental water samples. Based on publicly available TiLV genomic sequence data (Ahasan et al. 2020; Chaput et al. 2020; Debnath et al. 2020; Pulido et al. 2019; Subramaniam et al. 2019; Thawornwattana et al. 2021), we developed a new probe-based RT-qPCR assay targeting TiLV genomic segment 9 and applied to detect TiLV not only from fish tissues but also from environmental RNA (eRNA) concentrated from water samples. A simple iron flocculation method for concentration of TiLV from fish-rearing water samples coupled with our new RT-qPCR assay to detect and quantify TiLV eRNA was described in the present study.

## Materials & Methods

### Development of a new probe-based quantitative RT-qPCR method for TiLV

#### Primer & probe design and establishment of PCR conditions

A new hydrolysis probe-based RT-qPCR method was developed and optimized for detection and quantification of TiLV. Out of the 10 segments of the TiLV genome, segment 9 was reported to have relatively high identity (97.44 - 99.15%) among various TiLV isolates (Pulido et al. 2019). Primers and probe were thus designed based on conserved regions of TiLV genome segment 9 following multiple sequence alignments of all available sequences (n=25 or 27) retrieved from the GenBank database at NCBI as of June 2021 (Fig. S1). Primer Seg9-TaqMan-F (5’-CTA GAC AAT GTT TTC GAT CCA G-3’) had a 100% perfect match with all retrieved 27 sequences while primer Seg9-TaqMan-R (5’-TTC TGT GTC AGT AAT CTT GAC AG-3’) and probe (5’-6-FAM-TGC CGC CGC AGC ACA AGC TCC A-BHQ-1-3’) had one mismatch nucleotide from 25 and 27 available sequences, respectively (Fig. S1). The final composition of the optimized TiLV RT-qPCR 20 µL reaction consists of 1X master mix (qScript XLT 1-Step RT-qPCR ToughMix Low ROX buffer) (Quanta Bio), 1.5-2 µl (≤ 300 ng) of RNA template, 450 nM of each forward and reverse primers, and 150 nM of Seg9-TaqMan-Probe. Size of the amplified product is expected at 137 bp. Cycling conditions include a reverse transcription step at 50 °C for 10 min, then an initial denaturation step at 95 °C for 1 min followed by 40 cycles of 95 °C for 10 s and 58 °C for 30 s. RT-qPCR amplification was carried out using Bio-Rad CFX Connect Real-Time PCR machine. Positive control plasmid (pSeg9-351) was previously constructed by inserting a 351 bp-TiLV segment 9 open reading frame (ORF) into pGEM T-easy vector (Promega) as reported earlier (Thawornwattana et al. 2021).

#### Analytical specificity and sensitivity tests

Specificity of the Seg9-targeted RT-qPCR was tested with RNA extracted (150 ng/reaction) from clinically healthy tilapia, 15 common fish bacterial pathogens, and fish tissues infected with nervous necrosis virus (NNV), infectious spleen and kidney necrosis virus (ISKNV), or scale drop disease virus (SDDV) (Table S1). Detection limit of the method was investigated using 10-fold serial dilutions of pSeg9-351 plasmid template from 10^6^ to 1 copies/µL template. The assays were performed in duplicate. Calculation of viral copy numbers was performed using standard curves prepared by plotting the log_10_ of serial plasmid dilutions versus quantification cycle (Cq) values.

#### Diagnostic specificity and sensitivity of the assay

We assessed the Seg9 RT-qPCR assay against RNA extracted from 65 samples held in our laboratory. Forty-four samples originated from known TiLV outbreaks and 21 from known non-diseased samples (healthy tilapia). Diagnostic test results were obtained using semi-nested RT-PCR methods as described before (Dong et al. 2017a; Taengphu et al. 2020). Analytical specificity and sensitivity of the assay were calculated according to formulas described by Martin (1984) as:

- Sensitivity % = [number of true positive samples / (number of true positive samples + number of false negative samples)] × 100
- Specificity % = [number of true negative samples / (number of true negative samples + number of false positive samples)] × 100

### Optimization for viral concentration protocol

#### Virus preparation

Viral stock used in this study was isolated from TiLV-infected Nile tilapia using E-11 cell line,, a clone of the cell line SSN-1 derived from whole fry tissue of snakehead fish (Sigma-Aldrich cat no. 01110916-1VL). The virus was propagated as described in Dong et al. (2020). Briefly, 200 µL of TiLV stock (∼10^8^ copies/mL) was added into a 75 mL cell culture flask containing a monolayer of E-11 cell and 5 mL of L15 medium (Leibovitz), incubated at 25 °C for 5 days. The culture supernatant containing viral particles was collected after centrifugation at 15,000 x g for 10 min at 4 °C. The viral stock was kept in aliquots of 1 mL at −80 °C until used.

#### Iron flocculation

Viral concentration using iron flocculation method was performed using the protocol previously described by Kawato et al. (2016) with some modifications. Workflow of this method is illustrated in Fig. 1. Briefly, 100 µL (∼10^7^-10^8^ copies) of TiLV viral stock was added into 500 mL of sterile water that contained 1% marine salt and 36 µM ferric chloride. The suspension was stirred at room temperature for 1 h before being mechanically filtered through a 0.4-μm pore size polycarbonate filter (Advantec) with a vacuum pump connected to a filter holder KG-47 (Advantec) under < 15 psi pressure. The flocculate-trapped filters were either directly subjected to nucleic acid extraction or resuspended with oxalate-EDTA buffer (John et al. 2011) prior to nucleic acid extraction using Patho Gene-spin DNA/RNA extraction kit (iNtRON). Experiments were carried out in two to four replicates. Viral concentration, percentage (%) recovery and fold reduction of the virus copies were calculated from Cq values after flocculation compared to that of the starting viral stock.

**Figure 1:**
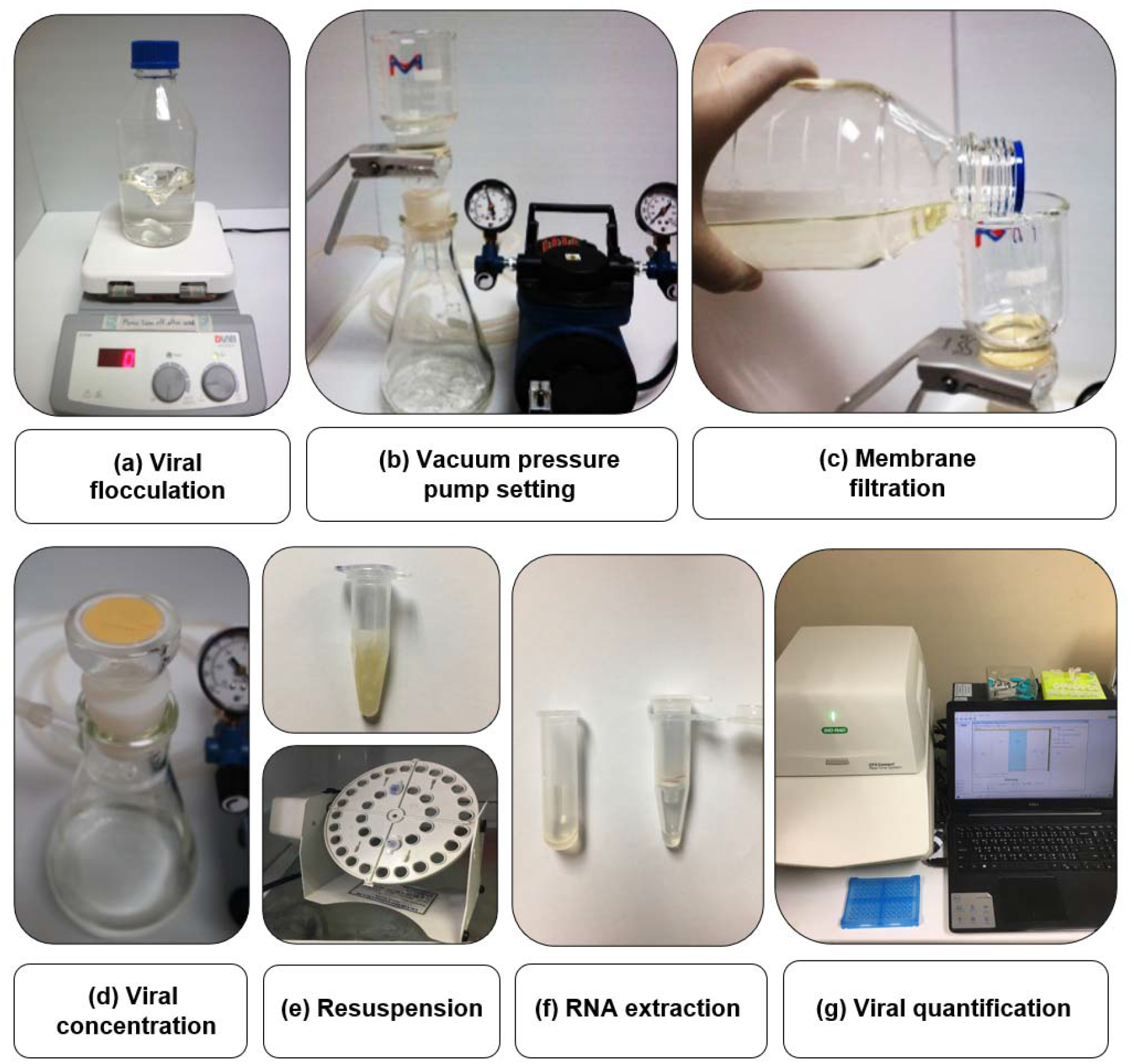
Workflow of TiLV flocculation, concentration and quantification used in this study. An iron flocculation method was used to concentrate viruses from water (a). The water suspension containing the virus was filtered through a 0.4-μm pore size polycarbonate membrane filter with a vacuum pressure pump (b-c). The flocculate-trapped filter (d) was then resuspended in oxalate-EDTA buffer (e) prior to nucleic acid extraction (f) and TiLV quantification (g).

### Detection of TiLV from fish and pond water sources during disease outbreaks

During 2020-2021, two disease outbreaks were reported to our laboratory. One occurred in an open-caged system (juvenile hybrid red tilapia, *Oreochromis* sp.) and the other in a closed hatchery system (earthen ponds, Nile tilapia, *O. niloticus*). The fish experienced abnormal mortalities with clinical symptoms of disease resembling those caused by TiLV, e.g. darkened body (Nile tilapia), pale color and reddish opercula (red hybrid tilapia), abdominal distension, and exophthalmia. In the first outbreak, we received fish specimens and water samples collected from four cages namely A, B, C and D with two-three fish and two bottles of 500 mL water samples from each cage. The samples were kept on ice during transportation and shipped to our laboratory within 24 h. In the latter outbreak, internal organs from both sick and healthy looking tilapia from different ponds as well as snails and sludge were collected and preserved in Trizol reagent (Invitrogen) by a hatchery veterinarian and sent to our laboratory. Water (500 mL/bottle) from fish ponds, reservoir, and sewage (outgoing waste water from ponds) was also collected from this hatchery.

Fish specimens were subjected to RNA extraction while water samples were centrifuged (5,000 x g for 5 min) to remove suspended matters before subjected to iron flocculation and subsequent nucleic acid extraction by Patho Gen-spin column kit. Viral detection and quantification were then performed to investigate the presence of TiLV by the established Seg 9 RT-qPCR assay described above. Plasmid template pSeg9-351 was used in a positive control reaction while nuclease-free water was used for negative control.

## Results

### A new probe-based RT-qPCR method for detection and quantification of TiLV

The Seg9 RT-qPCR method developed in this study had a detection limit (sensitivity) of 10 copies/µL template with mean Cq ± SD values of the detection limit at 38.24 ± 0.09 (Fig. 2a). Hence, samples with a Cq value ≥ 38.24 were considered TiLV negative or under the limit of this detection method. Amplification efficiency (E) of the established RT-qPCR was 94.0% with R^2^ of 0.998 (Fig. 2b). Analytical specificity test revealed that the method was highly specific to TiLV only since no amplifications were found when the method was assayed with RNA templates extracted from three other viruses, 15 bacterial species, and healthy tilapia (Fig. 2c, Table S1). The method had 100% diagnostic specificity and 100% diagnostic sensitivity when assayed with previously diagnosed TiLV infected and non-infected fish samples (n =65 with Cq value ranges 13.02 – 34.85) (Table 1).

**Table 1:** Diagnostic specificity and sensitivity of the Seg9 probe-based RT-qPCR method

| Test results | Diseased samples<br>(n=44) | Non-diseased samples<br>(n=21) |
| --- | --- | --- |
| Positive (+) | True positive<br>44 | False positive<br>0 |
| Negative (-) | False negative<br>0 | True negative<br>21 |
| Diagnostic sensitivity (%) | 100 |  |
| Diagnostic specificity (%) | 100 |  |

**Figure 2:**
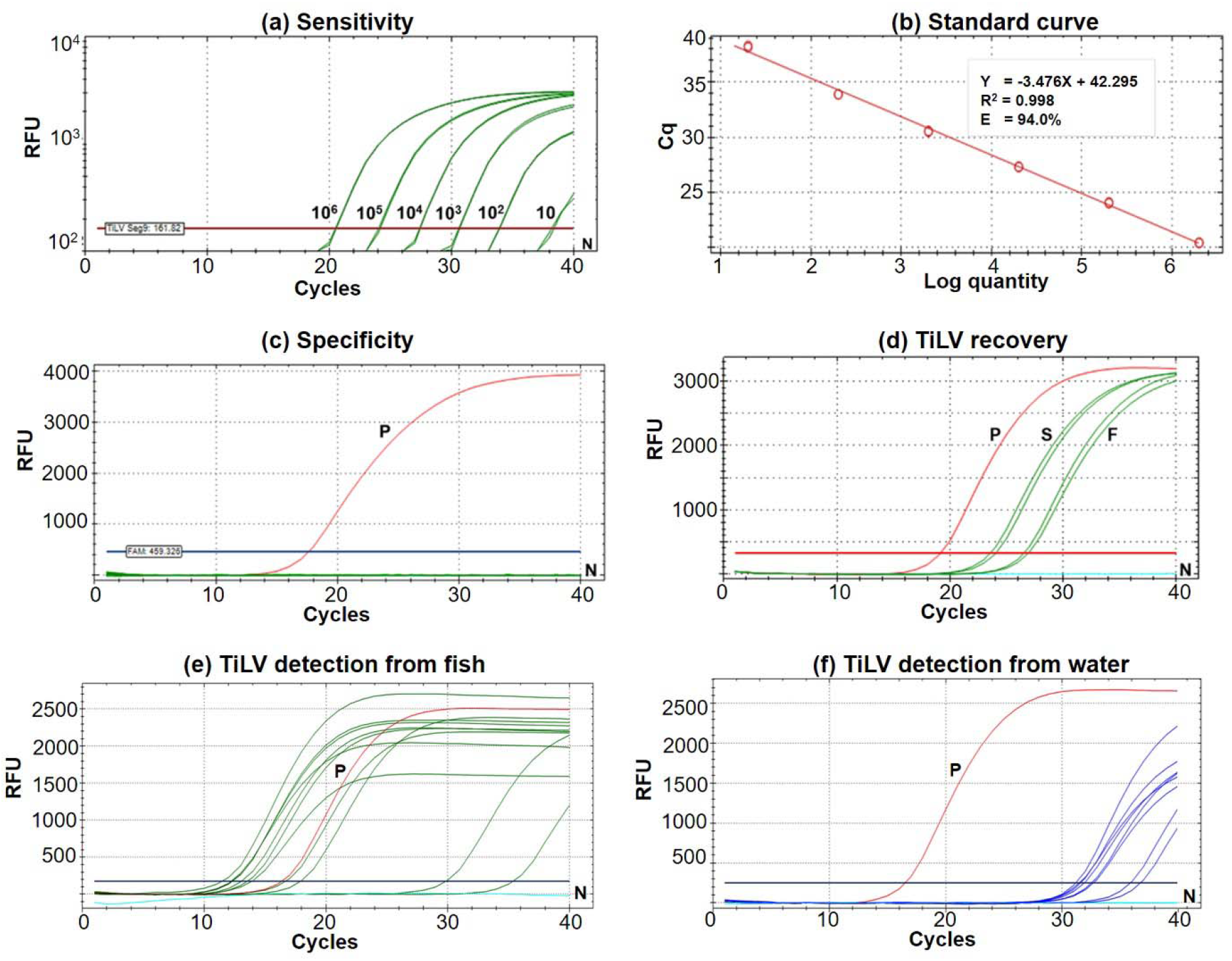
Performance of the newly established probe-based RT-qPCR detection of TiLV genomic segment 9. a) Analytical sensitivity assay determined using serial dilutions of plasmid DNA containing a 351-bp TiLV segment 9 insert. Amplification results were from two technical replicate tests. b) A standard curve was derived from the assays in (a) showing an amplification efficiency (E) of 94.0%. c) Analytical specificity test of the RT-qPCR protocol against RNAs extracted from common pathogens of fish and healthy looking tilapia as listed in Table S1. d) TiLV quantification from template extracted from stock virus (S) and flocculate-trapped filters (F) with resuspension step using two replicates. e) TiLV quantification from fish samples collected from an outbreak open cage. f) TiLV quantification from water samples collected from an outbreak open cage. P, positive control; N, no template control; RFU, relative fluorescence units.

### Conditions for viral concentration and percentage recovery

Percentage recovery of TiLV after iron flocculation but without suspension of the membrane filter in oxalate-EDTA buffer was only 2.04 ± 0.5% (n=2), which corresponded to a 50.55 ± 12.2-fold reduction in the viral concentration compared to the original viral stock (Table 2). This was significantly improved with an additional suspension step of the flocculate-trapped filters into oxalate-EDTA buffer prior to RNA extraction. The percentage recovery of TiLV increased to 16.11 ± 3.3% (n=4), which is equivalent to a 6.38 ± 1.1-fold reduction in viral concentration after iron flocculation (Table 2). Figure 2d showed representative results of viral quantification using Seg 9 RT-qPCR assays of TiLV from water after iron flocculation with the resuspension step.

**Table 2:** Percentage (%) recovery of viruses from water using different conditions

| Sample type | Before and after flocculation | Suspension step | Total viral copy number | % recovery | Fold reduction |
| --- | --- | --- | --- | --- | --- |
| Water spiked with TiLV culture | Before (viral stock) | | $3.92 \times 10^8$ | | |
| | After (Rep.1) | No | $9.34 \times 10^6$ | 2.38 | 41.93 |
| | After (Rep.2) | No | $6.62 \times 10^6$ | 1.69 | 59.18 |
|  | **Mean ± SD** |  |  | **$2.04 \pm 0.5$** | **$50.55 \pm 12.2$** |
| | Before (viral stock 1) | | $1.27 \times 10^8$ | | |
| | After (Rep.1) | Yes | $2.67 \times 10^7$ | 21.08 | 4.74 |
| | Before (viral stock 2) | | $3.21 \times 10^7$ | | |
| | After (Rep.2) | Yes | $4.67 \times 10^6$ | 14.55 | 6.87 |
| | Before (viral stock 3)* | | $4.16 \times 10^7$ | | |
| | After (Rep.3)* | Yes | $5.85 \times 10^6$ | 14.07 | 7.10 |
| | Before (viral stock 4)* | | $3.07 \times 10^7$ | | |
| | After (Rep.4)* | Yes | $4.52 \times 10^6$ | 14.74 | 6.78 |
|  | **Mean ± SD** |  |  | **$16.11 \pm 3.3$** | **$6.38 \pm 1.1$** |

Rep, replicate; \* denotes experiments where qPCR results were shown in Fig. 2d.

### Virus quantification from tilapia and different water sources during disease outbreaks

The results of TiLV detection and quantification from fish tissues and water samples are shown in Tables 3 and 4. In the first disease outbreak (open-cages), TiLV was detected from both fish and water samples from all four cages (A-D) (Table 3). Fish samples had Cq values ranging from 12.40 to 36.22, equivalent to 3.98 × 10^8^ to 5.6 × 10^1^ viral copies/150 ng RNA template, respectively (Table 3, Fig. 2e). Interestingly, eight water samples collected from four cages had a similar viral load ranging from 8.50 × 10^2^ to 3.40 × 10^4^ copies/L (Cq 31.19 - 36.76) (Table 3, Fig. 2f).

**Table 3:**
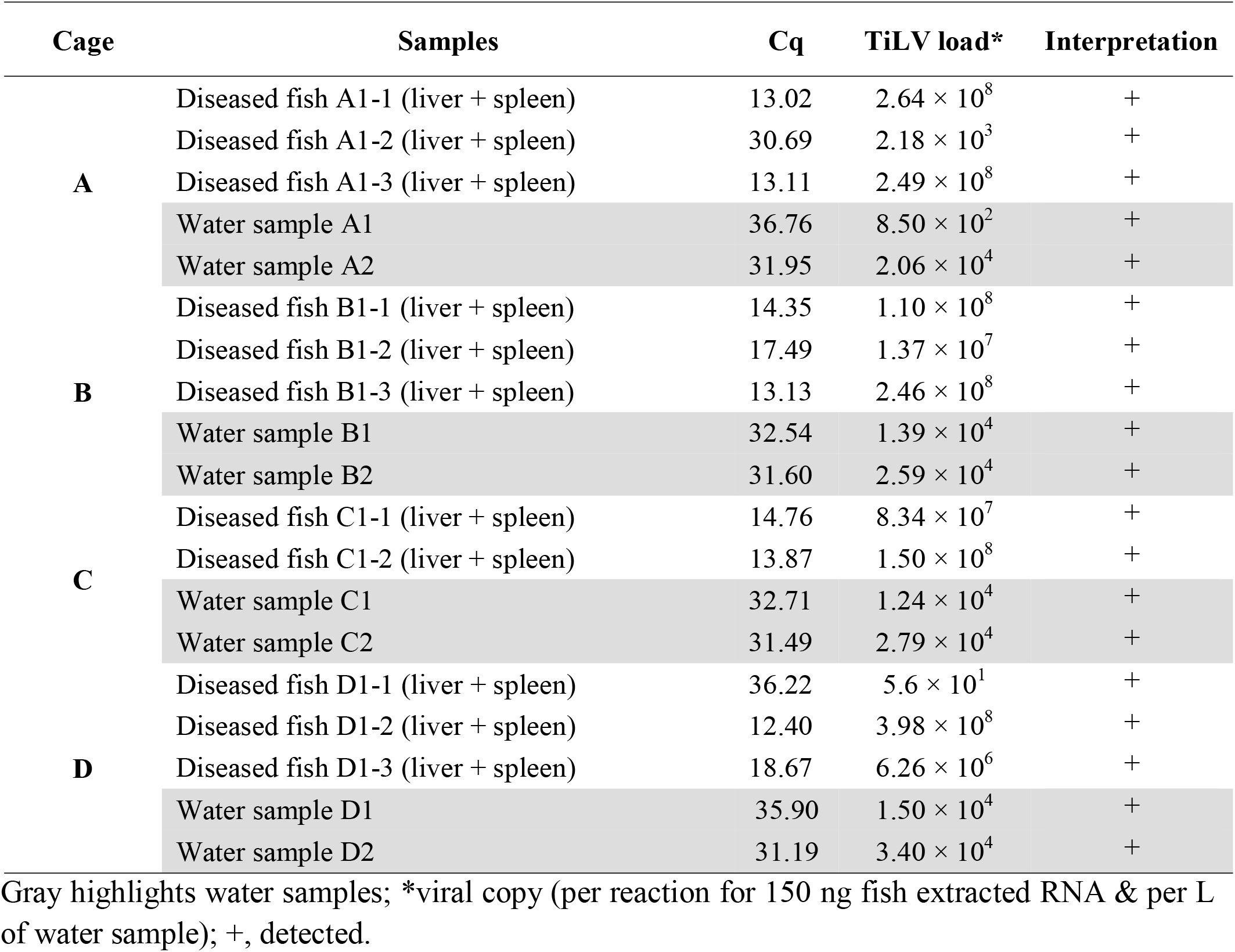
Quantification of TiLV from fish and water during an outbreak in open-cages

**Table 4:**
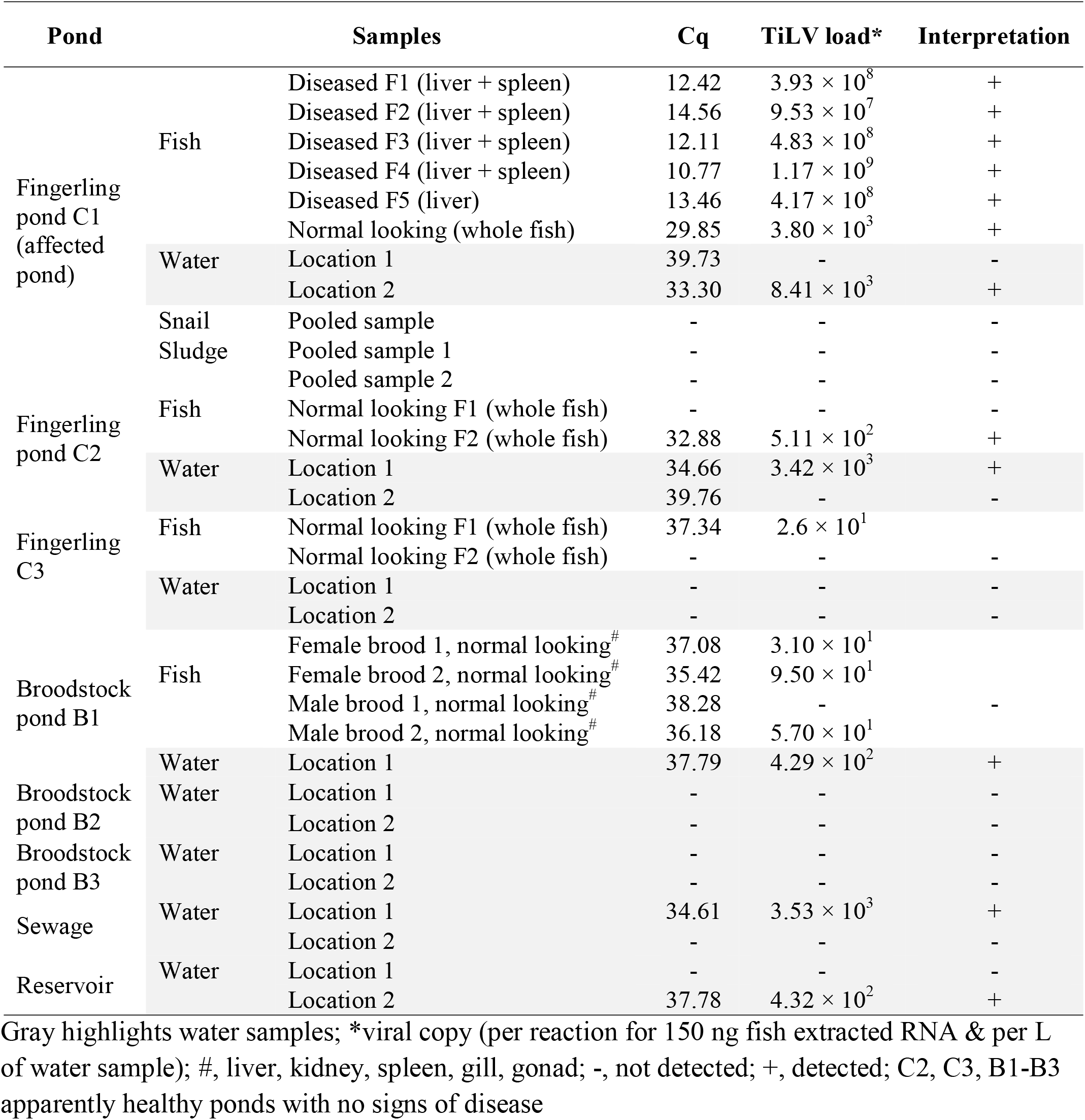
Quantification of TiLV from fish and pond water during an outbreak in earthen closed-ponds

In the second disease event (earthen ponds), samples were collected from eight ponds; one had unusually mortality, five showed no sign of disease, one was a sewage pond and one a reservoir pond (Table 4). In the affected fingerling pond C1, TiLV was detected from five diseased fish (9.53 × 10^7^ to 1.17 × 10^9^ copies/150 ng RNA template), one asymptomatic fish (3.80 × 10^3^ copies/150 ng RNA template), and water sample from one location (8.41 × 10^3^ copies/L) (Table 4). TiLV was undetectable from snail and sludge samples originating from pond C1. TiLV investigation from the remaining 7 other ponds revealed that TiLV was also detectable— but in relatively low viral loads from some asymptomatic fish (both fingerling and brood fish) and water from culture ponds as well as water from the reservoir and sewage ponds that were collected during the disease event (Table 4).

## Discussion

Methods to concentrate and recover viral particles from environmental water samples have been long applied in human health studies especially with waterborne diseases caused by enteric viruses (example review in Cashdollar & Wymer (2013); Haramoto et al. (2018)). It has later become an essential process for aquatic environment research (Jacquet et al. 2010). Several techniques have been used for viral concentration from aquatic environment, including coagulation/flocculation, filtration/ultrafiltration, and centrifugation/ultracentrifugation (Cashdollar & Wymer 2013; Ikner et al. 2012). Our present study employed an iron flocculation method which was initially described for virus removal from freshwater (Chang et al. 1958) and virus concentration from marine water (John et al. 2011). It was later adapted to detect and quantify two fish viruses: nervous necrosis virus (NNV) (an RNA virus) and red sea bream iridovirus (RSIV) (a DNA virus) that were experimentally spiked in fish-rearing water (Kawato et al. 2016; Nishi et al. 2016). The recovery rate was estimated by qPCR and yielded >50 and >80% for NNV and RSIV, respectively. In this study, while the recovery rate of TiLV (an RNA virus) from spiked-water was considerably lower (16.11 ± 3.3%), it is in a similar range of practical methods used for concentrating and detecting human viruses from water environments (Haramoto et al. 2018). For example, murine norovirus-1 (MNV-1) used as a viral model in viral concentration assay of human enteric viruses was recovered from spiked-water at 5.8–21.9% using the electronegative hydroxyapatite (HA)-filtration combined with polyethylene glycol (PEG) concentration method. The protocol was then used for detection of human noroviruses (NoV) and hepatitis A virus (HAV) in all water types (De Keuckelaere et al. 2013). More recently, researchers used porcine coronavirus (porcine epidemic diarrhea virus, PEDV) and mengovirus (MgV) as model viruses to concentrate severe acute respiratory syndrome coronavirus 2 (SARS-CoV-2) from water samples (Randazzo et al. 2020). By using an aluminum hydroxide adsorption-precipitation concentration method, PEDV and MgV spiked in water were recovered at 3.3-11.0%. The method can then be applied to detect SARS-CoV-2 RNA in untreated wastewater samples of ∼ 10^5.4^ genomic copies/L (Randazzo et al. 2020).

Despite a low recovery rate from water samples in this study, we confirmed the usefulness of the iron flocculation and RT-qPCR approach to concentrate and determine the concentration of TiLV from fish-rearing water and other water sources from two aquaculture production systems during disease outbreaks. The inherent nature of DNA and RNA viruses and their ability to survive outside their hosts may also contribute to those differences observed in recovery rates (Cashdollar & Wymer 2013; Pinon & Vialette 2018). Other viral concentration techniques using different coagulant/flocculant chemicals as well as more efficient RNA extraction methods should be tested for further improvement of TiLV recovery from water.

After the viral concentration and recovery process, downstream viral detection methods include cell culture methods, PCR-based assays, and viral metagenomics analysis (example review in Haramoto et al. (2018)). Here, we employed RT-qPCR technique for detection and quantification of TiLV, although the detected amounts did not represent the viral viability. Using all TiLV genomic sequences publicly available, we designed a new set of conserved primers and probe targeting the viral genomic segment 9. The newly established RT-qPCR protocol was highly specific to TiLV and did not cross-amplify RNA extracted from other common bacterial and viral aquatic pathogens. The method is very sensitive as it can detect as low as 10 viral copies per µL of template, >2,700 times more sensitive than previous probe-based RT-qPCR methods (Kembou Tsofack et al. 2017; Waiyamitra et al. 2018), reflecting high specificity of the newly designed primers and probe. Our RT-qPCR method has 100% diagnostic specificity and sensitivity in agreement with previous results (n=65) obtained using semi-nested RT-PCR protocols (Dong et al. 2017a; Taengphu et al. 2020). Increased number of sample sizes with diverse geographical sources may be required for further investigation. Most importantly, this new Seg 9 RT-qPCR assay was able to detect and quantify TiLV load from various types of field samples, including clinically sick fish, asymptomatic fish, and water samples, as opposed to other molecular diagnostic methods optimized solely for fish specimens.

The viral loads from water samples collected during the two disease events were approximately ∼ 10^3^ viral copies/L (earthen ponds) and ∼10^4^ viral copies/L (open-cages), but in reality, these concentrations might be significantly higher due to substantial losses during the concentration and recovery process. Higher viral loads observed in some of the water samples collected during the disease outbreak were probably due to active shedding of the virus from diseased fish into the environment, and might be an additional evidence of the waterborne transmission nature of TiLV reported previously (Eyngor et al. 2014; Yamkasem et al. 2019). Potential application for TiLV outbreak forecasting should be further investigated by experimental infection to monitor viral loads in water in relation to fish morbidity and mortality as previously described for other fish pathogens (Haramoto et al. 2007; Kawato et al. 2016; Minamoto et al. 2009; Nishi et al. 2016).

## Conclusions

In summary, the viral concentration method by iron flocculation used in concert with a newly developed probe-based RT-qPCR was not only successful for detection and quantification of TiLV from water in diseased pond/cages, but also from unaffected ponds, reservoir, and sewage water. This method, apart from its potential practical use for future monitoring programs of TiLV viral load in water samples from various culturing units, our approach could become useful to detect possible TiLV contamination from incoming and outgoing waste water as well as to test the systems after disinfection treatments. Such application will support health professionals and farmers to design appropriate biosecurity interventions to reduce the loss caused by TiLV in tilapia farms and hatcheries.

## Acknowledgements

This study was financially funded by the CGIAR Research Program on Fish Agri-Food Systems (FISH) led by WorldFish. The authors would like to thank K. Pimsannil, W. Meemetta and Ms. Thu Thao Mai for their skilled technical assistance.

## Supplementary data

**Figure S1.**
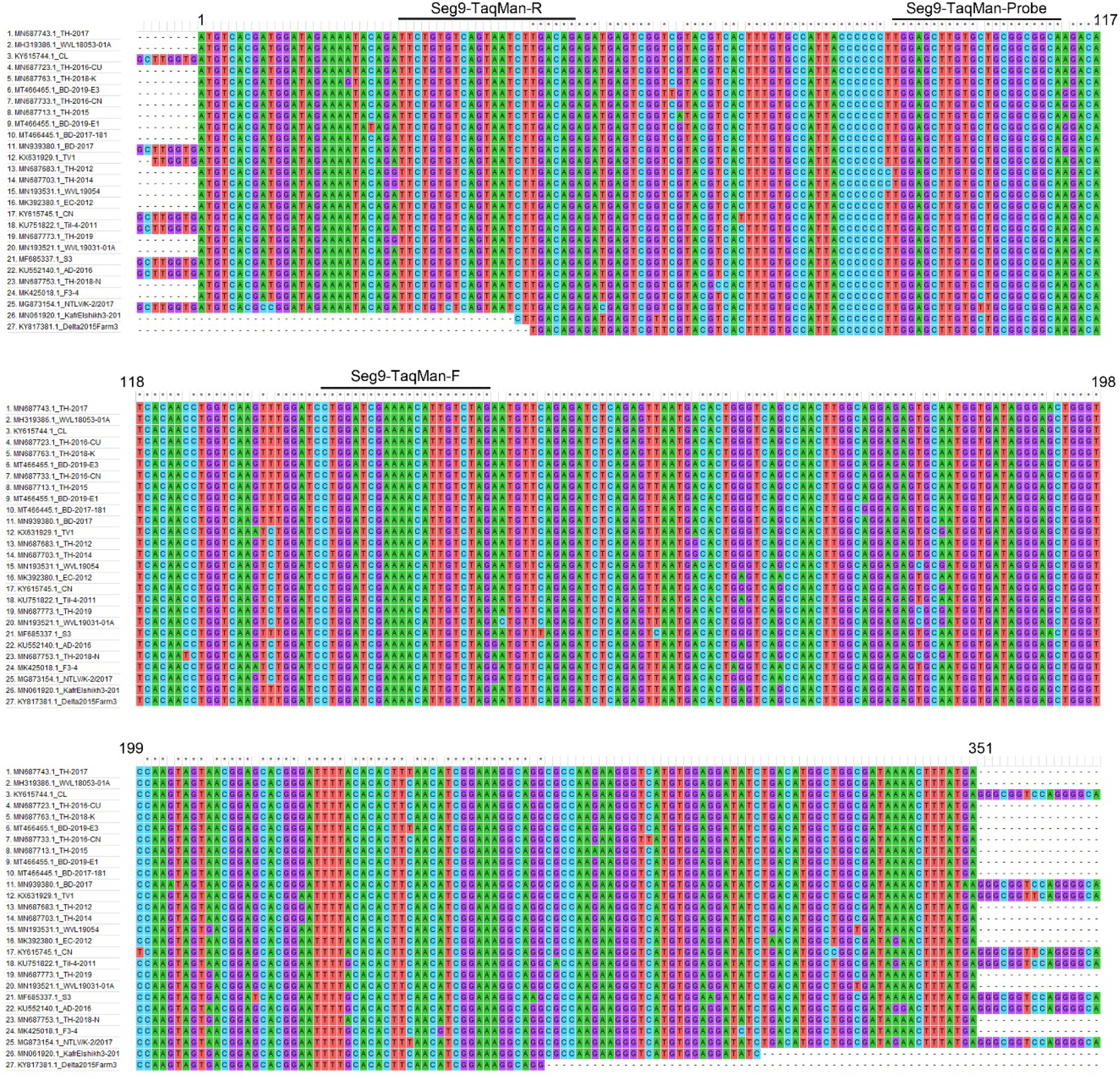
Nucleotide sequence alignments of TiLV segment 9 sequences (n=27) retrieved from the GenBank database at NCBI. Accession numbers and viral isolate names of all 27 sequences are shown on the left panel. Position of primers and probe used in the newly developed RT-qPCR assay are marked. Numbers denote nucleotide positions to the putative coding region.

**Table S1:** Sample used for evaluation of analytical specificity and sensitivity of the probe-based RT-qPCR method

| Samples | Host | Sample type | RT-qPCR result (Cq) | ND, not detectable |
| --- | --- | --- | --- | --- |
| Clinically healthy tilapia | NA | RNA | ND |  |
| NNV-infected tissue | Grouper | RNA | ND |  |
| ISKNV-infected tissue | Asian sea bass | RNA | ND |  |
| SDDV-infected tissue | Asian sea bass | RNA | ND |  |
| *Streptococcus agalactiae* | Nile tilapia | RNA | ND |  |
| *Streptococcus iniae* | Asian sea bass | RNA | ND |  |
| *Edwardsiella ictaluri* | Striped catfish | RNA | ND |  |
| *Edwardsiella tarda* | Nile tilapia | RNA | ND |  |
| *Flavobacterium columnare* | Asian sea bass | RNA | ND |  |
| *Francisella orientalis* | Hybrid red tilapia | RNA | ND |  |
| *Aeromonas hydrophila* | Tilapia | RNA | ND |  |
| *Aeromonas veronii* | Nile tilapia | RNA | ND |  |
| *Aeromonas dhakensis* | Hybrid red tilapia | RNA | ND |  |
| *Aeromonas caviae* | Nile tilapia | RNA | ND |  |
| *Aeromonas jandaei* | Nile tilapia | RNA | ND |  |
| *Plesiomonas shigelloides* | Nile tilapia | RNA | ND |  |
| *Chryseobacterium* sp. | Nile tilapia | RNA | ND |  |
| *Vogesella* sp. | Nile tilapia | RNA | ND |  |
| *Vibrio cholerae* | Nile tilapia | RNA | ND |  |

